# Pair housing does not alter incubation of craving, extinction, and reinstatement after heroin self-administration in female and male rats

**DOI:** 10.1101/2022.07.28.501777

**Authors:** Kelle E. Nett, Ryan T. LaLumiere

## Abstract

**Rationale:** Evidence suggests that single housing in rats acts as a chronic stressor, raising the possibilities that it contributes to measures of heroin craving and that pair housing ameliorates such measures.

**Objectives:** This study aimed to determine whether pair housing after heroin self-administration reduces the incubation of craving, extinction, and reinstatement of heroin seeking.

**Methods:** Single-housed female and male Sprague-Dawley rats underwent daily 6-h heroin self-administration, wherein active lever presses produced a heroin infusion paired with light/tone cues. One d after self-administration, rats underwent a baseline cued-seeking test wherein active lever presses only produced light/tone cues. Immediately following this cued-seeking test, rats were either pair-housed with a weight- and sex-matched naïve rat or remained single-housed for the rest of the study. For 14 d, rats remained in their homecages, after which they underwent a cued-seeking test to assess the incubation of craving compared to their baseline test. Rats then underwent extinction sessions followed by cue-induced and heroin-primed reinstatements.

**Results:** The findings reveal that pair-housed rats did not differ from single-housed rats in terms of the incubation of craving, extinction, or reinstatement of heroin seeking. Additionally, the results did not reveal any evidence of sex-based differences in the study.

**Conclusions:** The present work indicates that pair housing during the forced abstinence period does not alter measures of heroin craving/seeking. These findings suggest that the chronic stress of single housing specifically during forced abstinence does not contribute to the degree of such measures.

## Introduction

Evidence suggests that social support and interactions during recovery are beneficial for individuals with substance use disorder, including opioid use disorder, whereas social isolation is a risk factor for returning to drug use (Cohen and Wills, 1985; Havassy et al., 1991; Topor et al., 2011; Lookatch et al., 2019). Like humans, rats are highly social creatures, and evidence indicates that social isolation (e.g., single-housing conditions) alters behavioral and neurobiological measures compared to social housing (e.g., pair- or group-housing conditions) (Grippo et al., 2007a; Grippo et al., 2007c; Grippo et al., 2007b; Cacioppo et al., 2015; Sarkar and Kabbaj, 2016). For example, following 8 weeks of isolation, male rats display depression-like behavior and decreased spine density in the medial prefrontal cortex (Sarkar and Kabbaj, 2016). These effects likely result from the chronic stress that isolation produces in rodents, as evidence indicates that isolation in rats raises baseline corticosterone levels, increases adrenal weight, and decreases brain weight (Turner et al., 2014; Mastrogiovanni et al., 2021). Indeed, the alterations in medial prefrontal cortex with social isolation (Sarkar and Kabbaj, 2016) are similar to those observed with cocaine self-administration (Radley et al., 2015), and prior work indicates that this region is critical for promoting drug-seeking behavior, including heroin seeking (LaLumiere and Kalivas, 2008). Together with evidence indicating that stressors promote drug seeking (Mantsch et al., 2016), these findings raise the possibility that isolated housing contributes to measures of heroin craving and seeking.

Prior work examining aspects of substance use disorder in rodent models suggests that, under certain circumstances, social housing ameliorate some of the negative or maladaptive measures in these models. For example, socially-housed rats consume less drug, including opioids (Wolffgramm and Heyne, 1991; Raz and Berger, 2010; Westenbroek et al., 2013), and acquire heroin self-administration slower than single-housed rats (Westenbroek et al., 2019). One confounding factor of previous work examining isolated vs. social housing in drug seeking has been the concurrent use of environmental enrichment, such as running wheels, nesting materials, toys, and two or more cage mates (Nithianantharajah and Hannan, 2006; Kentner et al., 2021). When maintained with social housing as a component of environmental enrichment, rats extinguish heroin seeking faster, have decreased motivation to consume heroin, and have decreased drug-primed reinstatement of heroin seeking compared to isolated rats without environmental enrichment (Imperio et al., 2018). However, such studies do not isolate the social housing factor from the other enrichment factors.

Nonetheless, most opioid self-administration studies use single-housed rats, raising the possibility that isolation-induced stress contributes to commonly used measures of drug seeking. The increase in drug craving that happens over time during forced abstinence, known as the incubation of craving, occurs across drugs of abuse in humans and is readily observed in rats, allowing for increased translational relevance for manipulations that decrease drug craving (Venniro et al., 2016). In these experiments, rats typically remain isolated in their home cages after their last self-administration session for multiple weeks before cue-driven craving tests are given. Thus, isolation-induced stress may contribute to incubation of opioid craving and, conversely, pair housing rats after self-administration (i.e., during the incubation period) may reduce such craving. Other work indicates that social housing as a component of environmental enrichment given during the period of forced abstinence reduces measures of craving for natural rewards (Grimm et al., 2008) and cocaine (Thiel et al., 2012). Nonetheless, whether social housing following self-administration and during forced abstinence reduces the incubation of opioid craving is unknown.

To address this issue, single-housed female and male rats were trained to self-administer heroin during daily 6 h sessions. Following a baseline cued-seeking test, rats were pair-housed with a naïve rat of the same sex or remained single-housed for the rest of the study. Incubation of heroin craving was tested 14 d after self-administration. To determine whether such housing affected other measures of heroin seeking, rats then underwent extinction training and cued and heroin-primed reinstatement tests. Our results indicate that pair housing after self-administration did not reduce the incubation of craving in females or males and did not alter the extinction or reinstatement of heroin seeking.

## Methods

### Subjects

Female and male Sprague-Dawley rats (200-225 g and 225-250 g, respectively at the time of first surgery; Envigo; n=31) were used for this study. Of the total rats, 10 of them did not undergo any of the procedures described below and only served as the pair-housing partner. All rats were initially single-housed in a temperature-controlled environment under a 12 h light/dark cycle (lights on a 07:00) and allowed to acclimate to the vivarium at least 2 d before surgery. All procedures followed the National Institutes of Health guidelines for care of laboratory animals and were approved by the University of Iowa Institutional Animal Care and Use Committee.

### Surgery

Those rats that were to self-administer heroin (n = 21) were anesthetized with 5% isoflurane and maintained at 2-3% isoflurane. Meloxicam (2 mg/kg, s.c.) was administered as an analgesic before surgery, as well as 24 h after surgery. Rats also received sterile saline (3 ml, s.c.) after surgery for rehydration. Rats received catheter implants, wherein a rounded tip jugular vein catheter (SAI Infusion Technologies) was inserted 3.5 cm into the right jugular vein and sutured to the vein. Suture beads reinforced the tubing at the suture point. The opposite end of the catheter was externalized between the shoulder blades and connected to a harness with a 22-gauge guide cannula (PlasticsOne, Inc.), which was used for heroin delivery. Catheters were flushed 6 d per week with 0.1 ml of heparinized saline and glycerol to ensure catheter patency.

### Heroin Self-Administration

Self-administration training sessions were carried out 6 d per week in standard operant boxes, housed within sound-attenuating chambers (Med Associates, Fairfax, VT) and equipped with a central reward magazine flanked by two retractable levers. Cue lights were located directly above the levers, and a 4500 Hz Sonalert pure tone generator module was positioned above the right lever. A 6 W house light on the opposite wall of the operant chamber was illuminated throughout the training sessions.

Heroin (kindly provided by the National Institute on Drug Abuse) was dissolved in 0.9% sterile saline. A dose of 0.15 mg/kg/infusion heroin was used for the first 2 d of heroin self-administration, followed by 0.067 mg/kg/infusion heroin for all following days of self-administration. A lever press on the active (right) lever resulted in a 50 uL heroin infusion and a 5 s presentation of light and tone cues. A 20 s timeout period followed each lever press, during which additional active lever presses were recorded but had no scheduled consequence. Rats self-administered heroin for at least 12 d with > 10 infusions on 10 of the days and > 15 infusions on each of the final 3 d.

### Incubation of heroin craving after single or pair housing

After the above criteria were met and 1 d after the last self-administration session, rats underwent a 30 min cued-seeking test in which an active lever press resulted in light/tone cue presentation, but no heroin infusion. This test served as the baseline measure for incubation of craving (Test Day 1). Following the cued-seeking test, rats remained in their homecages for 14 d. Two groups characterized this phase. Some rats remained single-housed whereas others were pair-housed with a naive rat. Each naive rat had been previously single-housed, was weight-matched to its new cage mate, and was monitored for 1 h upon pairing for good socialization (i.e., no signs of aggression or fighting). Once paired, rats remained pair-housed until the completion of the experiment.

Following 14 d in the homecages, rats were returned to the operant chamber for a 1 h cued-seeking test (Test Day 14). Again, an active lever press resulted in light/tone cues but no heroin infusion. The first 30 min of this seeking test was used to assess incubation of heroin craving compared to the Test Day 1 baseline.

### Extinction and reinstatement of heroin seeking tests

One d after the last cued-seeking test, rats began daily 3 h extinction sessions, in which a lever press had no consequence. Rats continued extinction for at least 7 d, until the last 3 d had < 20 lever presses. Rats then underwent 3 h cue-induced and heroin-primed reinstatements in a counterbalanced manner, with at least 3 d of extinction with < 20 lever presses between each reinstatement. For cue-induced reinstatement, an active lever press resulted in light/tone cues. For heroin-primed reinstatement, rats received a small priming injection of heroin (0.25 mg/kg, s.c.) immediately prior to the reinstatement session, and active lever presses had no consequence.

### Statistical Analysis

All self-administration data were analyzed using a two-way repeated-measures ANOVA with day as the within-subject variable and housing (single vs pair) as the between-subject variable. For examining incubation of craving, active lever presses from Test Day 1 and the first 30 min of Test Day 14 were compared. To determine whether there were any important differences across the time course of active lever presses on Test Day 14, lever pressing was divided into four 15-minute bins and a two-way repeated-measures ANOVA was used, with bin as a within-subject factor and housing as a between-subject factor. For examining active lever presses during extinction, a two-way repeated-measures ANOVA with day as the within-subjects variable and housing as the between-subjects variable was used. For examining reinstatement, active lever pressing during the extinction baseline (an average of active lever presses 3 d immediately preceding reinstatement) and during the reinstatement test were the within-subject variables with housing as the between-subjects variable. The Greenhouse-Geisser correction was used if the assumption of sphericity was violated with the repeated-measures ANOVA. Sidak’s test was used for all multiple comparisons in *post-hoc* analysis. Although not fully powered by sex, each two-way repeated-measures ANOVA was also run separately for females and males as a preliminary analysis to identify any potential areas where differences may emerge and in accordance with NIH policy on sex as a biological variable. All data were analyzed in GraphPad Prism 9.0.0 (GraphPad Software, La Jolla, CA).

## Results

Fig. 1 shows the heroin self-administration data. Prior to pair or single housing during the forced abstinence period, the two groups did not display any significant differences during self-administration (Fig. 1b-d). The two-way repeated-measures ANOVA for active lever presses across self-administration (Fig. 1b) revealed no main effect of day (*F*_3.62, 69.01_ = 0.63, *p* = 0.62), no main effect of future housing (*F*_1, 19_ = 1.39, *p* = 0.25), and no interaction between housing and self-administration day (*F*_11, 209_ = 0.69, *p* = 0.75). The two-way repeated-measures ANOVA on infusions across self-administration (Fig. 1c) revealed a main effect of day (*F*_4.04, 76.74_ = 3.71, *p* < 0.01), no main effect of future housing (*F*_1, 19_ = 0.29, *p* = 0.60), and no interaction between housing and self-administration day (*F*_11, 209_ = 0.66, *p* = 0.78). The two-way repeated-measures ANOVA on mg/kg heroin across self-administration (Fig. 1d) revealed a main effect of day (*F*_3.96, 75.32_ = 3.65, *p* < 0.01), no main effect of future housing (*F*_1, 19_ = 0.27, *p* = 0.61), and no interaction between housing and self-administration day (*F*_11, 209_ = 0.68, *p* = 0.76). The same pattern of heroin self-administration was observed with females and males separately (Fig. 1e). Table 1 shows the statistical analyses for all three figures, split by sex, and likewise confirms that males and females show the same pattern of statistical results across the analyses. Collapsing across housing, a two-way repeated-measures ANOVA on mg/kg heroin across self-administration for female versus male rats (Fig. 1f) revealed a main effect of day (*F*_4.15, 78.90_ = 3.80, *p* < 0.01), no main effect of sex (*F*_1, 19_ = 1.35, *p* = 0.26), and no interaction between day and sex (*F*_11, 209_ = 0.85, *p* = 0.59). Together, these results suggest that there were no pre-existing differences between subsequently single- or pair-housed rats across self-administration, that infusions and mg/kg heroin increased for both groups across self-administration, and that female and male rats did not differ in heroin-taking measures.

**Table 1.** P-values from two-way repeated-measure ANOVA analyses in females and males separately

| Variable | Sex | Main Effects |  | Interaction |
| --- | --- | --- | --- | --- |
|  |  | Day/Time | Housing |  |
| Self-administration<br>(active lever presses) | F | $p = 0.55$ | $p = 0.21$ | $p = 0.99$ |
| | M | $p = 0.51$ | $p = 0.15$ | $p = 0.98$ |
| Self-administration<br>(infusions) | F | $p = 0.19$ | $p = 0.49$ | $p = 0.32$ |
| | M | $p = 0.10$ | $p = 0.47$ | $p = 0.97$ |
| Self-administration<br>(mg/kg heroin) | F | $p = 0.21$ | $p = 0.52$ | $p = 0.34$ |
| | M | $p = 0.11$ | $p = 0.47$ | $p = 0.97$ |
| Incubation of craving<br>(Test Day 1 vs Test Day 14) | F | **$*p = 0.03$** | $p = 0.35$ | $p = 0.61$ |
| | M | **$*p < 0.01$** | $p = 0.99$ | $p = 0.63$ |
| Incubation of craving<br>(timecourse of Test Day 14) | F | **$*p = 0.03$** | $p = 0.66$ | $p = 0.24$ |
| | M | **$*p = 0.02$** | $p = 0.87$ | $p = 0.24$ |
| Extinction<br>(active lever presses) | F | **$*p < 0.0001$** | $p = 0.53$ | $p = 0.52$ |
| | M | **$*p < 0.001$** | $p = 0.76$ | $p = 0.87$ |
| Cue-induced reinstatement<br>(active lever presses) | F | **$*p = 0.01$** | $p = 0.54$ | $p = 0.53$ |
| | M | **$*p = 0.001$** | $p = 0.87$ | $p = 0.90$ |
| Heroin-primed reinstatement<br>(active lever presses) | F | **$*p = 0.01$** | $p = 0.84$ | $p = 0.98$ |
| | M | **$*p = 0.01$** | $p = 0.47$ | $p = 0.51$ |

**Fig. 1.**
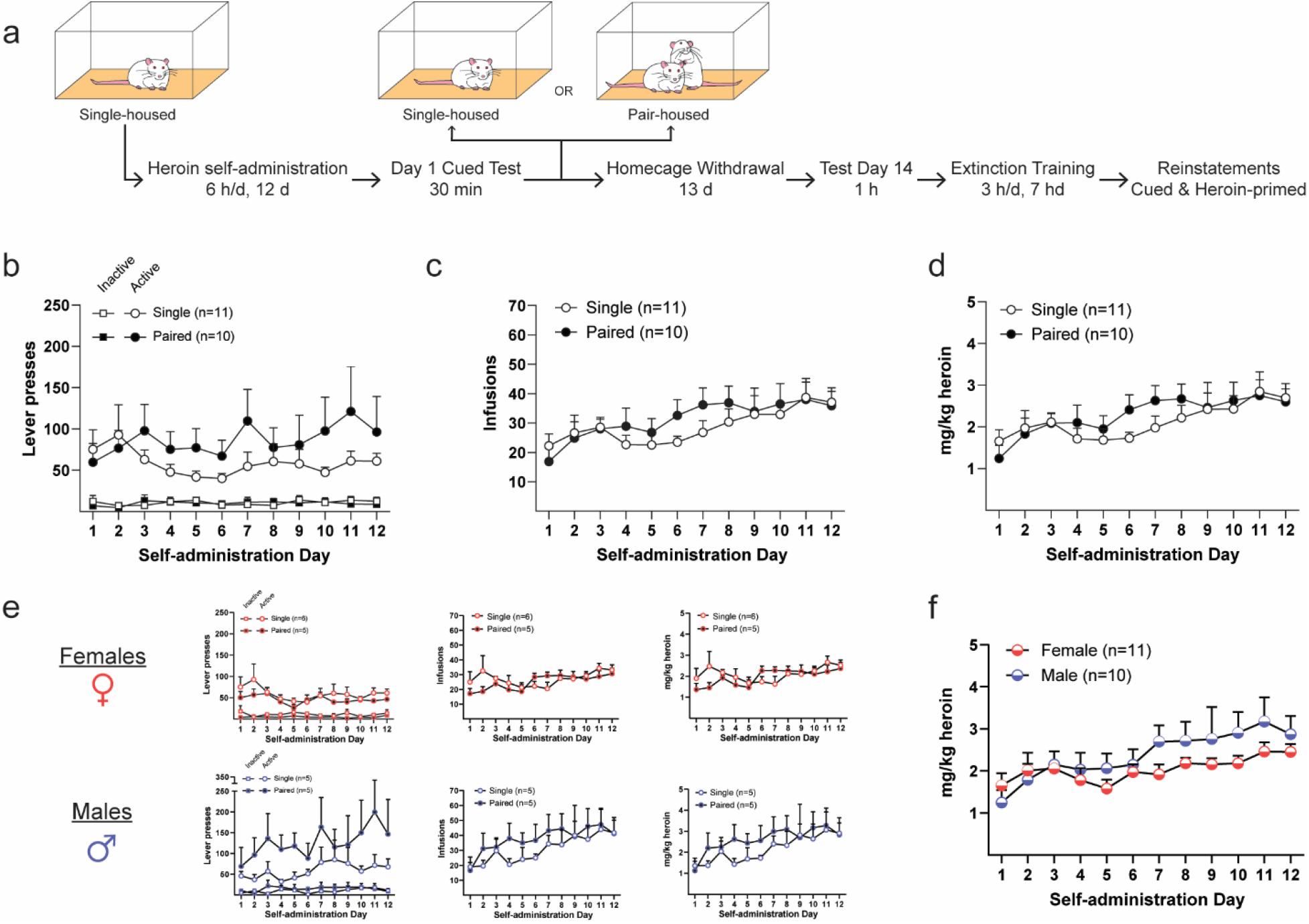
Experimental design and heroin self-administration. **a** Timeline of experiment. **b** Lever presses during heroin self-administration. Mean ± SEM of active (*right*) and inactive (*left*) lever presses across self-administration days. **c** Infusions during heroin self-administration displayed as mean ± SEM. **d** Mg/kg heroin self-administered. Mean ± SEM of total mg/kg heroin rats received each day of self-administration. **e** Lever presses (*left*), infusions (*middle*), and mg/kg heroin (*right*) during heroin self-administrations for females *(top, red*) and males (*bottom, blue*). **f** Mg/kg heroin self-administered between females and males. Mean ± SEM for all female and male rats, collapsed across pair-/single-housed conditions

Fig. 2 shows the results of pair versus single housing on the incubation of craving. The two-way repeated-measures ANOVA for active lever presses on Test Day 1 and 14 (Fig. 2a) revealed a main effect of test day (*F*_1, 19_ = 18.75, *p* < 0.001), no main effect of housing (*F*_1, 19_ = 0.20, *p* = 0.67), and no interaction between housing and test day (*F*_1, 19_ = 0.002, *p* = 0.96; Fig. 2a). To confirm the incubation of craving in both groups, *post-hoc* analyses revealed that both groups had significantly more lever presses on Test Day 14 compared to Test Day 1 (*p* < 0.05 in both cases). The two-way repeated-measures ANOVA for active lever presses during Test Day 14 across 15 min bins (Fig. 2b) revealed a main effect of bin (*F*_2.06, 39.16_ = 8.08, *p* < 0.01), no main effect of housing (*F*_1, 19_ = 0.06, *p* = 0.81; Fig. 2b), and an interaction between bin and housing (*F*_3, 57_ = 2.86, *p* = 0.05). *Post-hoc* analyses revealed no difference between housing group at any of the four timepoints (15 min, *p* = 0.84; 30 min, *p* = 0.20; 45 min, *p* = 0.95; 60 min, *p* = 0.88). The same pattern of active lever pressing during Test Day 14 was observed with both females and males (Fig. 2c; Table 1). These results suggest that pair housing during the incubation period did not affect the incubation of heroin craving.

**Fig. 2.**
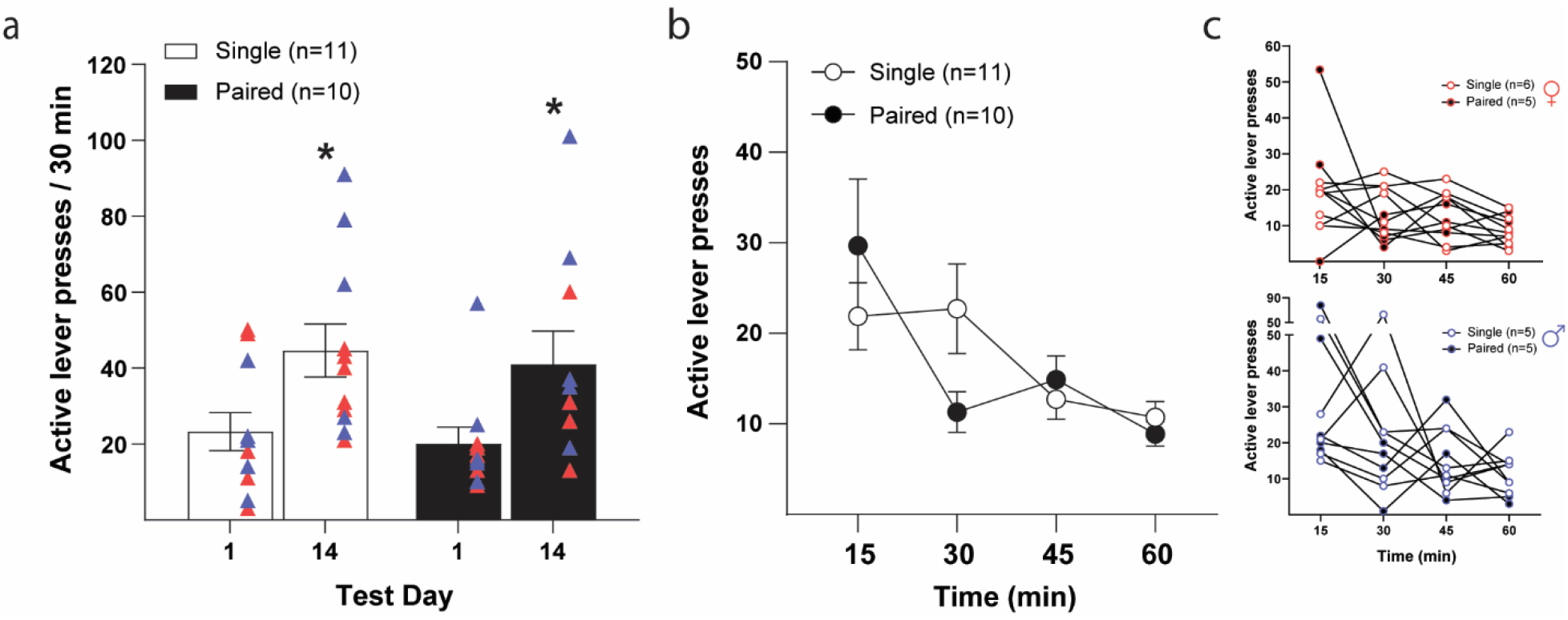
Incubation of heroin craving. **a** Active lever pressing on Test Day 1 and 14. Mean ± SEM of active lever presses during the 30 min Test Day 1 and from the first 30 min of Test Day 14. Triangles represent individual responses for females (*red*) and males (*blue*). *\*p* < 0.05 compared to Test Day 1. **b** Active lever presses during Test Day 14. Mean ± SEM of active lever presses every 15 min during the 1 h session. **c** Active lever presses as mean ± SEM during Test Day 14 between females (*top, red*) and males (*bottom, blue*)

Fig. 3 shows the active lever presses during extinction and cue-induced and heroin-primed reinstatement for the pair-versus single-housed rats. The two-way repeated-measures ANOVA of active lever presses during extinction (Fig. 3a) revealed a main effect of day (*F*_2.03, 38.61_ = 43.36, *p* < 0.0001), no main effect of housing (*F*_1, 19_ = 0.17, *p* = 0.68), and no interaction between day and housing (*F*_6, 114_ = 0.35, *p* = 0.91). A similar pattern of extinction lever pressing was observed in females and males separately (Fig. 3b; Table 1). The two-way repeated-measures ANOVA of active lever presses during cue-induced reinstatement (Fig. 3c) revealed a main effect of day (*F*_1,18_ = 33.08, *p* < 0.0001), no main effect of housing (*F*_1, 18_ = 0.43, *p* = 0.52), and no interaction between day and housing (*F*_1, 18_ = 0.43, *p* = 0.52). To confirm cue-induced reinstatement in both groups, *post-hoc* analyses revealed that active lever presses were significantly higher in both groups during cue-induced reinstatement compared to the extinction baseline (*p* < 0.01 in both cases). The two-way repeated-measures ANOVA of active lever presses during heroin-primed reinstatement (Fig. 3d) revealed a main effect of day (*F*_1,19_ = 22.32, *p* = 0.0001), no main effect of housing (*F*_1, 19_ = 0.17, *p* = 0.69), and no interaction between day and housing (*F*_1, 19_ = 0.22, *p* = 0.65). To confirm heroin-primed reinstatement in both groups, *post-hoc* analyses revealed that active lever presses were significantly higher in both groups during heroin-primed reinstatement test compared to extinction baseline (*p* < 0.05 in both cases). The same pattern of results for both cue-induced and heroin-primed reinstatement was observed in females and males separately (Fig. 3c-d, Table 1). These results reveal no effect of housing on the extinction of heroin seeking and cue-induced or heroin-primed reinstatement of heroin seeking.

**Fig 3.**
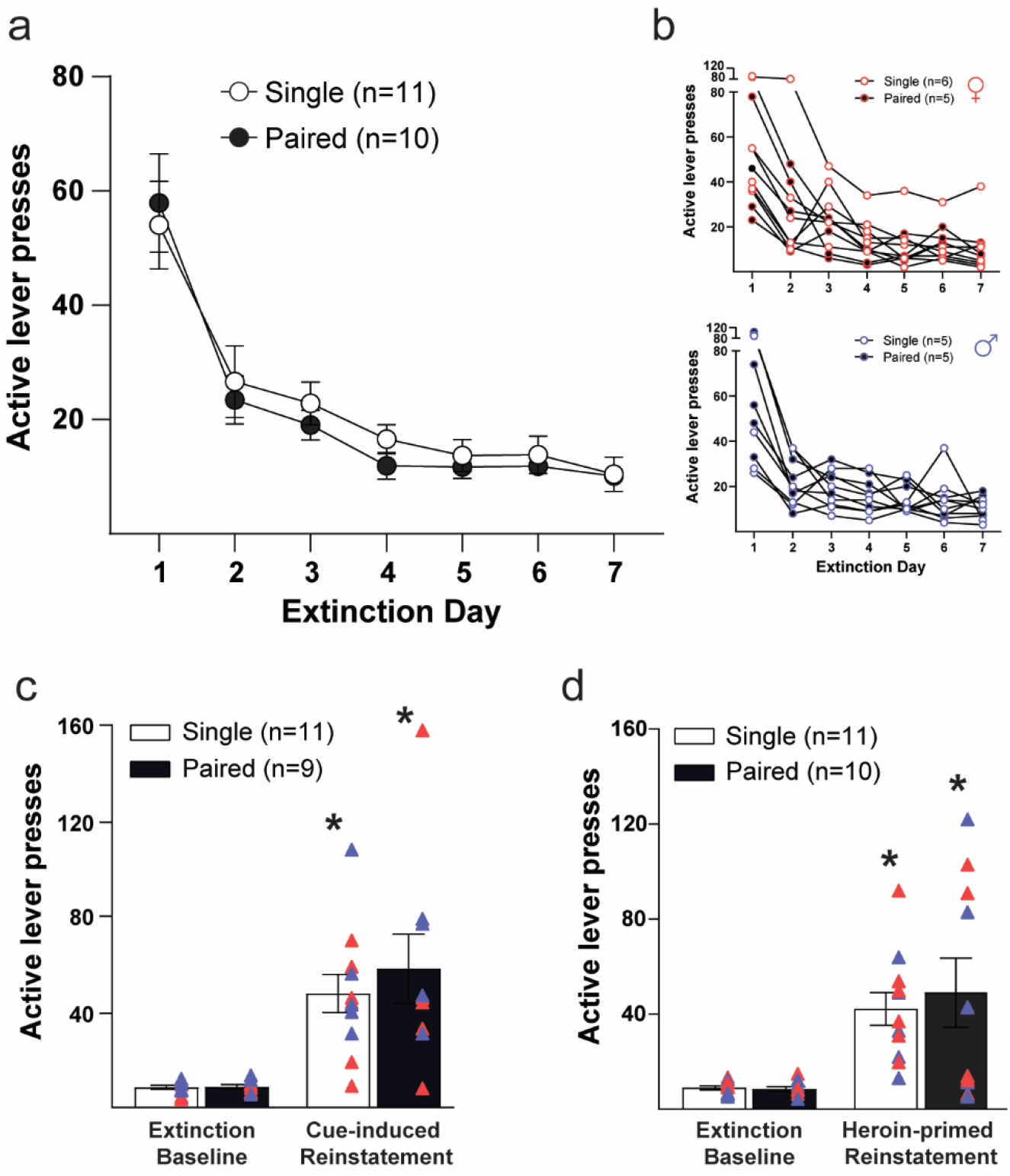
Extinction and reinstatement of heroin seeking. **a** Lever pressing throughout extinction. Mean ± SEM of active lever presses for the first 7 d of extinction. **b** Active lever pressing as mean ± SEM during extinction between females *(top, red*) and males (*bottom, blue*). **c** Cue-induced reinstatement of heroin seeking. Mean ± SEM of active lever presses from the last 3 d of extinction (extinction baseline) and from the cue-induced reinstatement session. **d** Heroin-primed reinstatement of heroin seeking. Mean ± SEM of active lever presses from the last 3 d of extinction (extinction baseline) and from the heroin-primed reinstatement session. Females, *red triangles*; Males, *blue triangles. *p* < 0.05 compared to extinction baseline

## Discussion

The present results indicate that pair housing during forced abstinence after heroin self-administration did not alter the incubation of heroin craving, the extinction of heroin seeking, or cue-induced or heroin-primed reinstatement. Moreover, of relevance for understanding the influence of sex on drug-related behavior, the findings did not reveal any sex differences in measures of heroin self-administration and craving/seeking or an interaction between sex and single vs. pair housing influences on such measures. Overall, the present study suggests that, at least with the procedures used herein, the ability of social interaction to attenuate measures of heroin craving and seeking is ineffective.

### Single housing during forced abstinence does not increase measures of heroin craving

Prior studies indicate that single housing during forced abstinence, compared to social housing with environmental enrichment, increases the incubation of sucrose and cocaine craving (Grimm et al., 2008; Thiel et al., 2012), as well as other measures of heroin seeking, such as the motivation to consume heroin and cue-induced heroin seeking (Imperio et al., 2018). However, the present work using only pair housing found no effect of such housing on various measures of heroin seeking, including incubation of craving. For social animals like rats, evidence indicates that prolonged isolation acts as a chronic stressor (Turner et al., 2014; Hofford et al., 2018; Mastrogiovanni et al., 2021), and stress influences drug seeking (Mantsch et al., 2016). Nonetheless, the present findings suggest that chronic social isolation, at least on its own, does not contribute to heroin craving/seeking.

There are two important distinctions between the present work and findings from Imperio et al. (2018): 1) the presence/absence of environmental enrichment and 2) the timing of the behavioral manipulations. In the study by Imperio et al. (2018), rats were socially housed in conjunction with environmental enrichment, whereas, in the present study, rats did not receive environmental enrichment when pair housed. It is possible that environmental enrichment more effectively alters heroin seeking compared to social housing. Indeed, for single-housed rats, there is evidence that environmental enrichment alone reduces cue-induced heroin seeking (Galaj et al., 2016; Imperio et al., 2018; Barrera et al., 2021), prevents heroin conditioned place preference (Galaj et al., 2016), and decreases compulsive cocaine and heroin seeking (Peck et al., 2015; Ewing and Ranaldi, 2018). Alternatively, social housing may only have effects as an interaction between housing and environmental enrichment.

Moreover, Imperio et al. (2018) kept their rats in their housing conditions beginning with self-administration, whereas the housing manipulations in the present study began during the forced abstinence period (i.e., after self-administration). Thus, it is possible that social housing must be present from self-administration onward for it to influence subsequent drug seeking. Prior studies indicate that single-housed rats, compared to socially housed rats *without environmental enrichment*, acquire heroin self-administration faster (Bozarth et al., 1989), consume more morphine (Raz and Berger, 2010), and reacquire heroin conditioned place preference faster (Turner et al., 2014).

Such findings suggest that social housing during self-administration itself protects against some effects of heroin taking, though the neurobiological mechanism for this is unclear. One such mechanism may involve oxytocin, as evidence suggests that chronic opioid use alters endogenous oxytocin signaling (You et al., 2000; Zanos et al., 2014), and enhancing oxytocin has been shown to reverse drug-induced neuroadaptations underlying tolerance and withdrawal (King et al., 2020) and decrease heroin self-administration (Kovacs et al., 1985). Oxytocin also enhances the social buffering of fear, or the ability of a conspecific to attenuate behavioral and autonomic measures of conditioned fear (Kiyokawa et al., 2007; Kiyokawa et al., 2014; Morozov and Ito, 2019; Peen et al., 2021). Taken together, these findings raise the possibility that oxytocin-enhancing manipulations, such as the pair housing used in the present study, has limited efficacy following chronic opioid use, but if provided throughout self-administration, acts as a social buffer of heroin craving by increasing oxytocin signaling, potentially preventing alterations in endogenous oxytocin signaling that occur from chronic opioid use.

Unlike many studies examining pair housing with drug seeking, the current study used a 6-h “long-access” self-administration procedure. To our knowledge, only one study examining the effects of environmental enrichment on heroin seeking used long-access (≥ 6 h) heroin self-administration and found that environmental enrichment during forced abstinence reduces the incubation of heroin craving (Sikora et al., 2018). However, that work did not address the effect of social housing itself, as all rats were housed in triads. Indeed, future work examining the effects of social housing should consider whether greater access to the drug via long-access self-administration overwhelms the effects of social housing to ameliorate different drug-related behaviors.

Notably, the present study observed a significant interaction between housing condition and time during the 1 h Test Day 14 cued seeking test (Fig. 2b). However, *post hoc* analyses revealed no significant differences at any individual time point. It is likely that the interaction reflects the inherent bin-by-bin variability that occurs across a 1-h cued-seeking test rather than a meaningful finding, particularly given that all other measures did not differ based upon housing conditions. Overall, evidence from the present study observed no differences between measures of heroin seeking and incubation of heroin craving for rats that were single-versus pair-housed, suggesting that such measures are not influenced by isolation during forced abstinence. However, it is unclear to what degree methodological details (i.e., heroin dosage, length of self-administration, timing of pair housing, etc.,) influence these measures. It is likely that subtle differences in self-administration procedures, including length of session, drug dosage, and total days of self-administration, contribute to measures of drug craving and seeking, potentially obscuring findings from manipulations with a smaller effect size.

### Female and male rats self-administer heroin similarly

Although not the focus of the present study, females and males were used in the experiments and no sex differences in heroin taking, craving, or seeking measures were observed. There were also no apparent interactions between housing condition and sex, though the groups were not fully powered to detect such differences. Critically, female and male rats undergoing long-access (6 h) heroin self-administration did not differ in total mg/kg heroin consumed. Some studies report sex differences in certain measures of opioid taking (Lynch and Carroll, 1999; Carroll et al., 2001; Becker and Koob, 2016; George et al., 2021). For example, George et al. (2021) found that female rats displayed increased responding and intake across heroin doses following 6-h heroin self-administration. However, not all studies report such differences (Lynch and Carroll, 1999; D’Ottavio et al., 2022). D’Ottavio et al. (2022) found that females had increased incubation of heroin craving compared to males, but only after intermittent access, and not continuous access, heroin self-administration. Nonetheless, a recent comprehensive review surveying the field found that studies examining measures of opioid craving in rodents largely did not reveal sex differences (Nicolas et al., 2022). Taken together with the present findings, it appears that, when sex differences are observed in opioid taking, craving, and/or seeking, they likely depend upon particular methodological parameters rather than reflecting widespread and robust differences between females and males.

### Conclusion

The present findings add to our understanding of psychosocial factors and their ability to influence opioid craving and seeking in rodents. The absence of differences between single- and pair-housed rats indicates that, during forced abstinence from heroin, housing condition does not contribute to the incubation of heroin craving or other measures of heroin seeking.

## Notes

**Funding** This work was supported by DA048055 (RTL).

**Conflict of Interest** Authors have no conflicts of interest to declare.

### Competing Interest Statement

The authors have declared no competing interest.

## References

Barrera ED, Loughlin L, Greenberger S, Ewing S, Hachimine P, Ranaldi R (2021) Environmental enrichment reduces heroin seeking following incubation of craving in both male and female rats. Drug Alcohol Depend 226:108852.

Becker JB, Koob GF (2016) Sex Differences in Animal Models: Focus on Addiction. Pharmacol Rev 68:242–263.

Bozarth MA, Murray A, Wise RA (1989) Influence of housing conditions on the acquisition of intravenous heroin and cocaine self-administration in rats. Pharmacol Biochem Behav 33:903–907.

Cacioppo JT, Cacioppo S, Capitanio JP, Cole SW (2015) The neuroendocrinology of social isolation. Annu Rev Psychol 66:733–767.

Carroll ME, Campbell UC, Heideman P (2001) Ketoconazole suppresses food restriction-induced increases in heroin self-administration in rats: sex differences. Exp Clin Psychopharmacol 9:307–316.

Cohen S, Wills TA (1985) Stress, social support, and the buffering hypothesis. Psychol Bull 98:310–357.

D’Ottavio G, Reverte I, Ragozzino D, Meringolo M, Milella MS, Boix F, Venniro M, Badiani A, Caprioli D (2022) Increased heroin intake and relapse vulnerability in intermittent relative to continuous self-administration: Sex differences in rats. Br J Pharmacol.

Ewing S, Ranaldi R (2018) Environmental enrichment facilitates cocaine abstinence in an animal conflict model. Pharmacol Biochem Behav 166:35–41.

Galaj E, Manuszak M, Ranaldi R (2016) Environmental enrichment as a potential intervention for heroin seeking. Drug Alcohol Depend 163:195–201.

George BE, Barth SH, Kuiper LB, Holleran KM, Lacy RT, Raab-Graham KF, Jones SR (2021) Enhanced heroin self-administration and distinct dopamine adaptations in female rats. Neuropsychopharmacology 46:1724–1733.

Grimm JW, Osincup D, Wells B, Manaois M, Fyall A, Buse C, Harkness JH (2008) Environmental enrichment attenuates cue-induced reinstatement of sucrose seeking in rats. Behav Pharmacol 19:777–785.

Grippo AJ, Cushing BS, Carter CS (2007a) Depression-like behavior and stressor-induced neuroendocrine activation in female prairie voles exposed to chronic social isolation. Psychosom Med 69:149–157.

Grippo AJ, Lamb DG, Carter CS, Porges SW (2007b) Social isolation disrupts autonomic regulation of the heart and influences negative affective behaviors. Biol Psychiatry 62:1162–1170.

Grippo AJ, Gerena D, Huang J, Kumar N, Shah M, Ughreja R, Carter CS (2007c) Social isolation induces behavioral and neuroendocrine disturbances relevant to depression in female and male prairie voles. Psychoneuroendocrinology 32:966–980.

Havassy BE, Hall SM, Wasserman DA (1991) Social support and relapse: commonalities among alcoholics, opiate users, and cigarette smokers. Addict Behav 16:235–246.

Hofford RS, Prendergast MA, Bardo MT (2018) Modified single prolonged stress reduces cocaine self-administration during acquisition regardless of rearing environment. Behav Brain Res 338:143–152.

Imperio CG, McFalls AJ, Hadad N, Blanco-Berdugo L, Masser DR, Colechio EM, Coffey AA, Bixler GV, Stanford DR, Vrana KE, Grigson PS, Freeman WM (2018) Exposure to environmental enrichment attenuates addiction-like behavior and alters molecular effects of heroin self-administration in rats. Neuropharmacology 139:26–40.

Kentner AC, Speno AV, Doucette J, Roderick RC (2021) The Contribution of Environmental Enrichment to Phenotypic Variation in Mice and Rats. eNeuro 8.

King CE, Gano A, Becker HC (2020) The role of oxytocin in alcohol and drug abuse. Brain Res 1736:146761.

Kiyokawa Y, Takeuchi Y, Mori Y (2007) Two types of social buffering differentially mitigate conditioned fear responses. Eur J Neurosci 26:3606–3613.

Kiyokawa Y, Hiroshima S, Takeuchi Y, Mori Y (2014) Social buffering reduces male rats’ behavioral and corticosterone responses to a conditioned stimulus. Horm Behav 65:114–118.

Kovacs GL, Borthaiser Z, Telegdy G (1985) Oxytocin reduces intravenous heroin self-administration in heroin-tolerant rats. Life Sci 37:17–26.

LaLumiere RT, Kalivas PW (2008) Glutamate release in the nucleus accumbens core is necessary for heroin seeking. J Neurosci 28:3170–3177.

Lookatch SJ, Wimberly AS, McKay JR (2019) Effects of Social Support and 12-Step Involvement on Recovery among People in Continuing Care for Cocaine Dependence. Subst Use Misuse 54:2144–2155.

Lynch WJ, Carroll ME (1999) Sex differences in the acquisition of intravenously self-administered cocaine and heroin in rats. Psychopharmacology (Berl) 144:77–82.

Mantsch JR, Baker DA, Funk D, Le AD, Shaham Y (2016) Stress-Induced Reinstatement of Drug Seeking: 20 Years of Progress. Neuropsychopharmacology 41:335–356.

Mastrogiovanni NA, Wheeler AK, Clemens KJ (2021) Social isolation enhances cued-reinstatement of sucrose and nicotine seeking, but this is reversed by a return to social housing. Sci Rep 11:2422.

Morozov A, Ito W (2019) Social modulation of fear: Facilitation vs buffering. Genes Brain Behav 18:e12491.

Nicolas C, Zlebnik NE, Farokhnia M, Leggio L, Ikemoto S, Shaham Y (2022) Sex Differences in Opioid and Psychostimulant Craving and Relapse: A Critical Review. Pharmacol Rev 74:119–140.

Nithianantharajah J, Hannan AJ (2006) Enriched environments, experience-dependent plasticity and disorders of the nervous system. Nat Rev Neurosci 7:697–709.

Peck JA, Galaj E, Eshak S, Newman KL, Ranaldi R (2015) Environmental enrichment induces early heroin abstinence in an animal conflict model. Pharmacol Biochem Behav 138:20–25.

Peen NF, Duque-Wilckens N, Trainor BC (2021) Convergent neuroendocrine mechanisms of social buffering and stress contagion. Horm Behav 129:104933.

Radley JJ, Anderson RM, Cosme CV, Glanz RM, Miller MC, Romig-Martin SA, LaLumiere RT (2015) The Contingency of Cocaine Administration Accounts for Structural and Functional Medial Prefrontal Deficits and Increased Adrenocortical Activation. J Neurosci 35:11897–11910.

Raz S, Berger BD (2010) Social isolation increases morphine intake: behavioral and psychopharmacological aspects. Behav Pharmacol 21:39–46.

Sarkar A, Kabbaj M (2016) Sex Differences in Effects of Ketamine on Behavior, Spine Density, and Synaptic Proteins in Socially Isolated Rats. Biol Psychiatry 80:448–456.

Sikora M, Nicolas C, Istin M, Jaafari N, Thiriet N, Solinas M (2018) Generalization of effects of environmental enrichment on seeking for different classes of drugs of abuse. Behav Brain Res 341:109–113.

Thiel KJ, Painter MR, Pentkowski NS, Mitroi D, Crawford CA, Neisewander JL (2012) Environmental enrichment counters cocaine abstinence-induced stress and brain reactivity to cocaine cues but fails to prevent the incubation effect. Addict Biol 17:365–377.

Topor A, Borg M, Di Girolamo S, Davidson L (2011) Not just an individual journey: social aspects of recovery. Int J Soc Psychiatry 57:90–99.

Turner PV, Sunohara-Neilson J, Ovari J, Healy A, Leri F (2014) Effects of single compared with pair housing on hypothalamic-pituitary-adrenal axis activity and low-dose heroin place conditioning in adult male Sprague-Dawley rats. J Am Assoc Lab Anim Sci 53:161–167.

Venniro M, Caprioli D, Shaham Y (2016) Animal models of drug relapse and craving: From drug priming-induced reinstatement to incubation of craving after voluntary abstinence. Prog Brain Res 224:25–52.

Westenbroek C, Perry AN, Becker JB (2013) Pair housing differentially affects motivation to self-administer cocaine in male and female rats. Behav Brain Res 252:68–71.

Westenbroek C, Perry AN, Jagannathan L, Becker JB (2019) Effect of social housing and oxytocin on the motivation to self-administer methamphetamine in female rats. Physiol Behav 203:10–17.

Wolffgramm J, Heyne A (1991) Social behavior, dominance, and social deprivation of rats determine drug choice. Pharmacol Biochem Behav 38:389–399.

You ZD, Li JH, Song CY, Wang CH, Lu CL (2000) Chronic morphine treatment inhibits oxytocin synthesis in rats. Neuroreport 11:3113–3116.

Zanos P, Georgiou P, Wright SR, Hourani SM, Kitchen I, Winsky-Sommerer R, Bailey A (2014) The oxytocin analogue carbetocin prevents emotional impairment and stress-induced reinstatement of opioid-seeking in morphine-abstinent mice. Neuropsychopharmacology 39:855–865.

